# Epigenetic clock and methylation studies in the rhesus macaque

**DOI:** 10.1101/2020.09.21.307108

**Authors:** Steve Horvath, Joseph A. Zoller, Amin Haghani, Anna J. Jasinska, Ken Raj, Charles E. Breeze, Jason Ernst, Julie A. Mattison

## Abstract

Methylation levels at specific CpG positions in the genome have been used to develop accurate estimators of chronological age in humans, mice, and other species. Although epigenetic clocks are generally species-specific, the principles underpinning them appear to be conserved at least across the mammalian class. This is exemplified by the successful development of epigenetic clocks for mice and several other mammalian species. Here, we describe epigenetic clocks for the rhesus macaque (*Macaca mulatta*), the most widely used nonhuman primate in biological research. Using a custom methylation array (HorvathMammalMethylChip40), we profiled n=281 tissue samples (blood, skin, adipose, kidney, liver, lung, muscle, and cerebral cortex). From these data, we generated five epigenetic clocks for macaques. These clocks differ with regards to applicability to different tissue types (pan-tissue, blood, skin), species (macaque only or both humans and macaques), and measure of age (chronological age versus relative age). Additionally, the age-based human-macaque clock exhibits a high age correlation (R=0.89) with the vervet monkey (*Chlorocebus sabaeus*), another Old World species. Four CpGs within the *KLF14* promoter were consistently altered with age in four tissues (adipose, blood, cerebral cortex, skin). It is expected that the macaque clocks will reveal an epigenetic aging rate associated with a host of health conditions and thus lend themselves for identifying and validating anti-aging interventions.

## INTRODUCTION

The rising costs of healthcare have fueled the growing need to address the root cause of most diseases and health conditions. Age is, without doubt, the strongest correlative factor across a wide range of pathologies. As such, investigations into the mechanisms and causes of aging, as well as interventions that might ameliorate its effects, hold great promise for improving health. To meet this end, animal models that closely recapitulate human aging are essential. Rhesus macaques (*Macaca mulatta*) are the most widely used nonhuman primate in biomedical research and share over 92% DNA sequence homology with humans (*1*). They have an average lifespan in captivity of approximately 27 years, maximal lifespan of 42 years, and experience aging processes that are very similar to humans. With these features, the rhesus macaque presents as an excellent subject for the understanding of aging in humans and also other closely-related primate species (*2, 3*). Despite these attractive features, the employment of rhesus macaques in such research remains modest. This is due to both the prohibitive cost of maintaining a colony and the relatively long lifespan of these primates (*4*). These challenges, however, can be effectively addressed if accurate and robust biomarkers of biological age are established. Such biomarkers would change the experimental endpoint from longevity (measure of time from birth to death), to biological age (the measure of health and fitness). The application of biomarkers will greatly reduce the duration and cost of primate studies, while generating a much more meaningful understanding of why we age and provide the means to evaluate anti-aging interventions.

We report here, the development of DNA methylation-based biomarkers of age, known as epigenetic clocks. The development of such clocks was inspired by the vision that combining methylation levels of multiple CpGs that change with age would produce an accurate age estimator (reviewed in (*5–7*)). This notion was made possible by the technical advancement of array platforms that provide accurate quantitative measurements of methylation for thousands of specific CpGs in the genome and subsequently furthered the development of several human epigenetic clocks. Human and mouse pan-tissue DNA methylation (DNAm) age estimators exhibited important characteristics for aging studies, namely application to all sources of DNA (from sorted cells, tissues, and organs) and across the entire age spectrum (from prenatal tissue to centenarians) (*5, 8–10*). A substantial body of literature report that human epigenetic clocks capture many aspects of biological age (*5*). As such, the discrepancy between DNAm age and chronological age (termed as “epigenetic age acceleration”) is predictive of all-cause mortality in humans even after adjusting for a variety of known risk factors (*11–13*). Several age related conditions are also associated with epigenetic age acceleration, including, but not limited to, cognitive and physical functioning (*14*), centenarian status (*13, 15*), Down syndrome (*16*), HIV infection (*17*), obesity (*18*).

Recently, mouse epigenetic clocks were successfully applied to evaluate and confirm benchmark longevity interventions such as calorie restriction and ablation of growth hormone receptor (*9, 10, 19–22*). These observations strongly suggest that the capture of biological age by epigenetic clocks is not the preserve of human clocks but is a feature that applies several mammalian species. Since age-related DNA methylation change appears to be evolutionarily conserved, accurate age estimators, such as those developed for humans, may be applied across species. Although the human pan-tissue clock can indeed be applied to chimpanzee DNA methylation profiles, its performance with profiles of other animals decline as a result of evolutionary genome sequence divergence (*8*). Here we describe the development and performance of several epigenetic clocks for rhesus macaques, two of which are dual-species clocks that apply both to humans and rhesus macaques.

## RESULTS

### DNA methylation data

All rhesus macaque DNA methylation profiles were generated on a custom methylation array (HorvathMammalMethylChip40) that measures the methylation level of 36,000 CpGs with flanking DNA sequences that are conserved across the mammalian class. We obtained 281 DNA methylation profiles from 8 different tissues of rhesus macaque (*Macaca mulatta*) with ages that ranged from 1.8 years to 42 years (**Table 1**). An unsupervised hierarchical analysis clustered the methylation profiles by tissue type (**Supplementary Figure 1**). DNA methylation-based age estimators (epigenetic clocks) were developed using data from n=281 tissues, of which the most numerous were blood (N=199) and skin (N=51). Postmortem tissues (omental adipose, brain cortex, kidney, liver, lung, and skeletal muscle) were also available, but from fewer than 7 animals (**Table 1**). To generate dual-species epigenetic clocks that apply to both human and rhesus macaque, n=850 human tissue samples that were similarly profiled on HorvathMammalMethylChip40 were added to the rhesus macaque training set (**Methods**).

**Table 1.** Description of biological materials from which DNA methylation profiles were derived. N=Total number of tissues. Number of females. Age: mean, minimum and maximum in units of years.

| Tissue | N | No.Female | Mean.Age | Min.Age | Max.Age |
| --- | --- | --- | --- | --- | --- |
| Adipose | 5 | 2 | 31.3 | 23.5 | 42 |
| Blood | 199 | 71 | 17.2 | 1.79 | 42 |
| Cortex | 6 | 3 | 29 | 17.2 | 42 |
| Kidney | 4 | 1 | 28.7 | 23.5 | 33.4 |
| Liver | 5 | 4 | 25.1 | 17.2 | 42 |
| Lung | 6 | 3 | 29 | 17.2 | 42 |
| Muscle | 5 | 2 | 30.1 | 17.2 | 42 |
| Skin | 51 | 13 | 18.8 | 7.61 | 42 |

### Epigenetic clocks

From these datasets, we generated five epigenetic clocks for macaques. These clocks differ with regards to applicability to different tissue types (pan-tissue, blood, skin), species (macaque only or both humans and macaques), and measure of age (chronological age versus relative age). As indicated by their names, pan-tissue clocks apply to all tissues, while the other clocks are developed for specific tissues/organs (blood, skin). The macaque pan-tissue clock was trained on all available tissues and applies only to rhesus macaques. The two human-macaque pan-tissue clocks, on the other hand, were derived from DNA methylation profiles from both species and are distinct from each other based on the unit of age that is employed. One estimates *chronological age* (in units of years), while the other estimates *relative* age, which is the ratio of chronological age to maximum lifespan, with values between 0 and 1. This ratio allows alignment and biologically meaningful comparison between species with very different lifespan (rhesus macaque and human), which is not afforded by mere measurement of chronological age. The maximum recorded lifespans for rhesus macaques and humans are 42 years and 122.5 years, respectively, according to the updated version of the *anAge* data base (*23*), thus there is an approximate 3:1 age ratio.

To arrive at unbiased estimates of the rhesus macaque pan-tissue clock, we carried out cross-validation analysis of the training data, followed by evaluation with an independent data set from another nonhuman primate species (vervet monkey). The cross-validation study reports unbiased estimates of the age correlation R (defined as Pearson correlation between the DNAm age estimate and chronological age) as well as the median absolute error.

The resulting macaque pan-tissue clock is highly accurate in age-estimation across tissues (R=0.95, median absolute error 1.4 years, **Figure 1A**) and in individual types (R>=0.93, **Supplementary Figure 2C-I**), except for adipose tissue (R=0.73, **Supplementary Figure 2B**) for which only n=5 samples were available. The human-rhesus macaque clock for age is highly accurate when both species are analyzed together (R=0.98, **Figure 1D**), with a slight reduction when the analysis is restricted to rhesus macaque tissues (R=0.95, **Figure 1E**).

**Figure 1:**
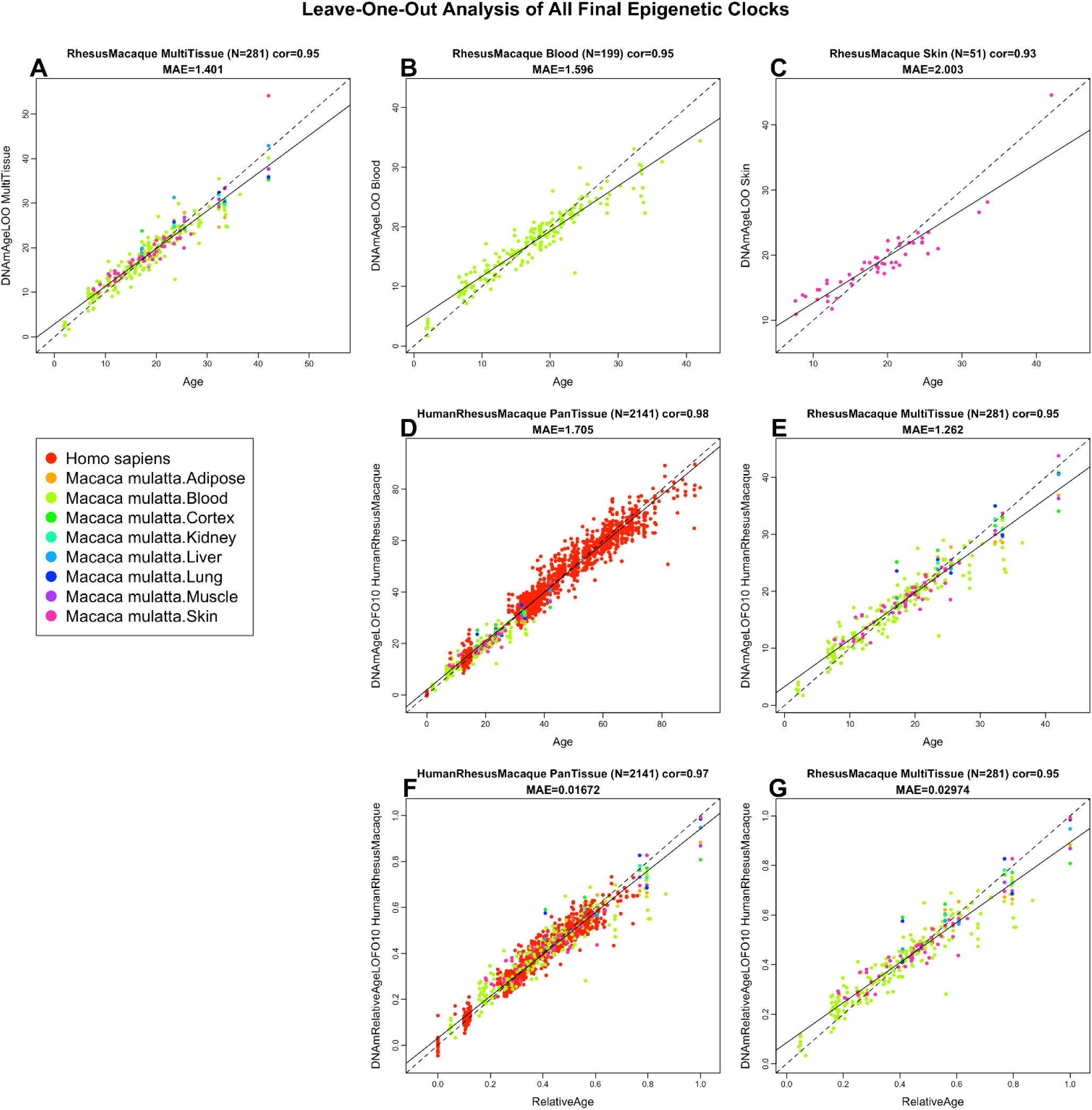
Cross-validation study of epigenetic clocks for rhesus macaques and humans. A-C) Three epigenetic clocks that apply only to macaques. Leave-one-sample-out estimate of DNA methylation age (y-axis, in units of years) versus chronological age for A) all available macaque tissues, B) blood, C) skin. Ten-old cross validation analysis of the human-macaque monkey clocks for D,E) chronological age and F,G) relative age, respectively. D,F) Human samples are colored in red and macaque samples are colored by macaque tissue type, and analogous in E,G) but restricted to macaque samples (colored by macaque tissue type). Each panel reports the sample size (in parenthesis), correlation coefficient, median absolute error (MAE).

Similarly, the human-rhesus macaque clock for *relative age* exhibits high correlation regardless of whether the analysis is done with samples from both species (R=0.97, **Figure 1F**) or with only rhesus macaque samples (R=0.95, **Figure 1G**). The employment of relative age circumvents the inevitable unequal distribution of data at the opposite ends of the age range when chronological age of species with very different lifespans are measured using a single formula. A cross validation analysis reveals that both human-macaque clocks lead to high accuracy (R>=0.97) in human blood and skin samples (**Figure 2**).

**Figure 2.**
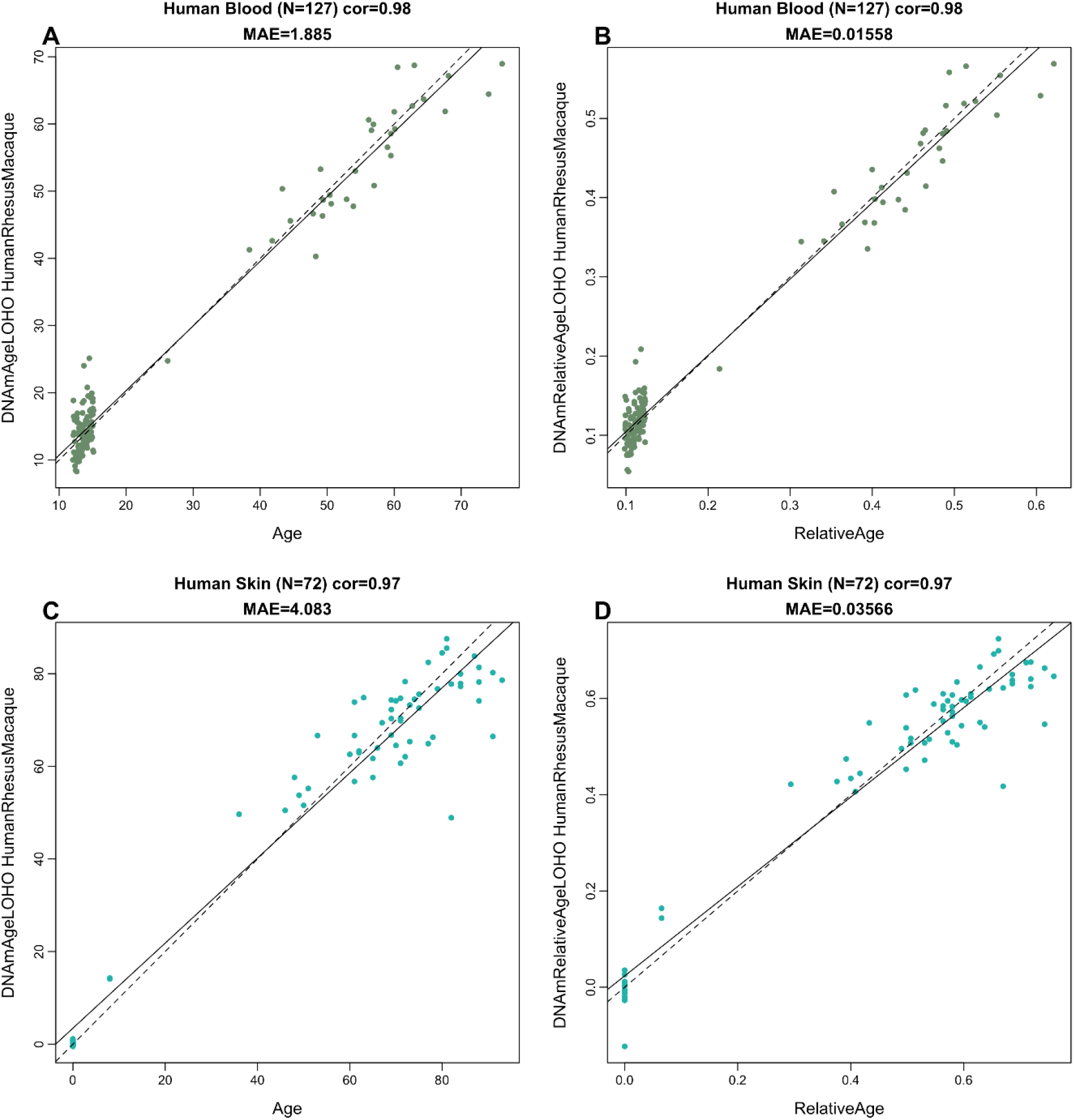
Human-macaque clocks applied to select human tissues. Leave-one-human sample-out (LOHO) cross fold cross validation estimates of the human-macaque clock for A,C) chronological age and C,D) relative age, respectively. A,B) Human blood samples. C,D) Human skin samples. Each panel reports the sample size (in parenthesis), correlation coefficient, median absolute error (MAE).

### Cross-species performance of the rhesus macaque pan-tissue clock

To determine the extent by which the rhesus macaque epigenetic clock can be applied to another primate, we used it to estimate the age of numerous tissues (blood, brain cortex, and liver) of the vervet monkey (*Chlorocebus sabaeus*), which is another Old World monkey separated 12.5 million years ago from the macaques. Despite this, we observed high correlations between the chronological age of vervets and their predicted age based on the macaque pan-tissue clock: R=0.96 in vervet blood, R=0.92 in vervet cortex, and R=0.98 in vervet liver (**Figure 3A-D**). It is worth noting that the comparison of correlation coefficients between different tissues is not straightforward as these values are dependent on the age distribution of the samples that are evaluated (e.g. minimum and maximum age) and also on the sample size, albeit to a lesser extent. While the correlations are nevertheless impressively high, the level of concordance between chronological age and estimated age are less so, in particular with cortex, which exhibited an off set of 9 years (**Figure 3B**). Nevertheless, there is reasonably good concordance between chronological age of vervets and the estimated age of their blood (median error 1.7 years, **Figure 3B**) and liver (median error 3.5 years, **Figure 3C**) by the macaque pan-tissue clock.

**Figure 3.**
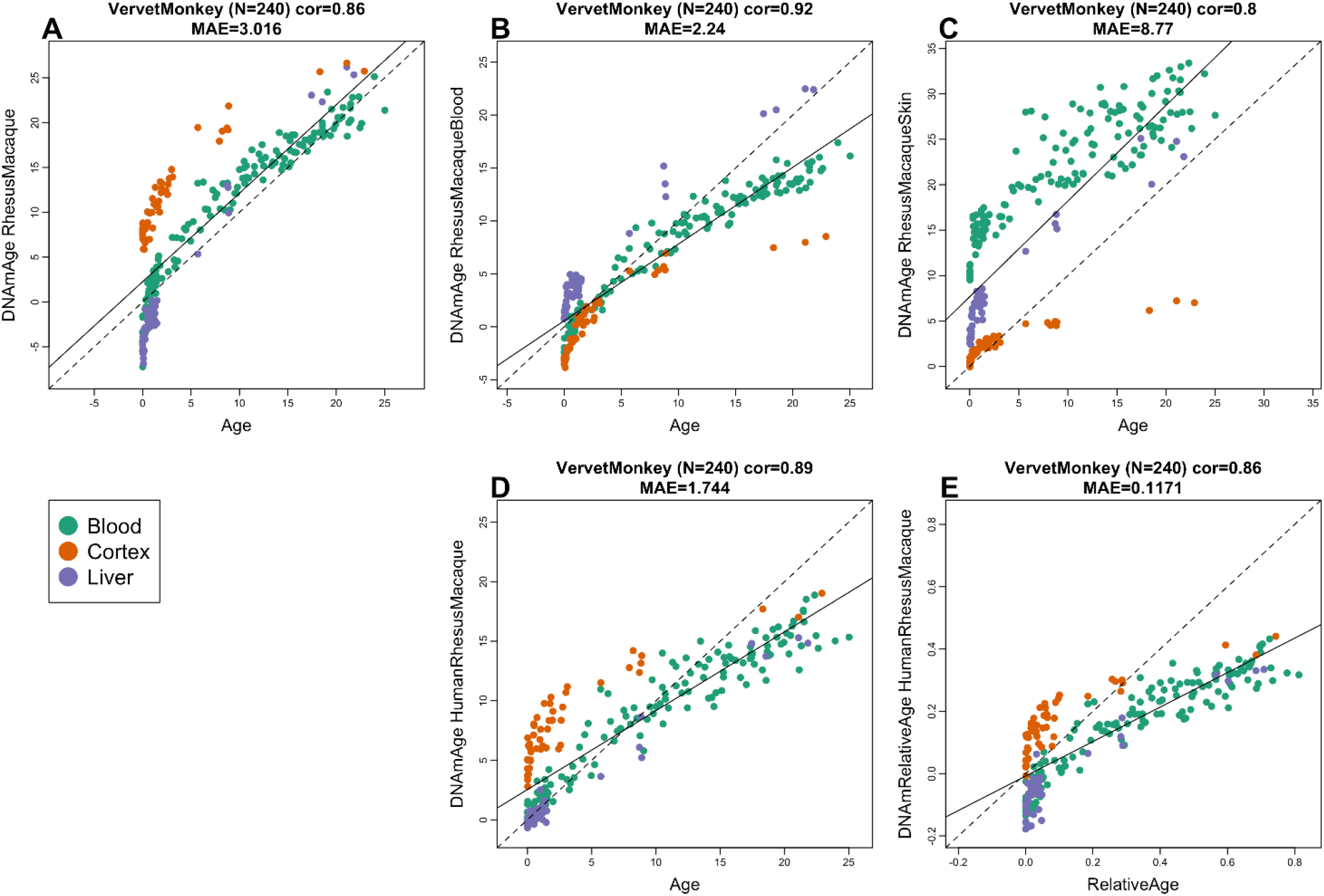
Macaque clocks applied to tissues from vervet monkey (*Chlorocebus sabaeus*). Each dot corresponds to a tissue sample from vervet monkeys. Each dot is colored by tissue type: blood (green), cerebral cortex (red), liver (purple). Chronological age of the vervet specimens (x-axis) versus the DNAm age estimate of the A) pan-tissue macaque clock, B) blood macaque clocks, C) skin macaque clock, D) human-macaque clock for chronological age, E) human-macaque clock for relative age. Each panel reports the sample size (in parenthesis), correlation coefficient, median absolute error (MAE).

### Epigenome-Wide Association Studies (EWAS) of chronological age in rhesus Macaque

In total, 36,733 probes from HorvathMammalMethylChip40 could be mapped to specific loci in rhesus Macaque (*Macaca mulatta*.Mmul_10.100) genome. These loci are located proximal to 6154 genes. It is expected that findings resulting from the use of these clocks can be extrapolated to humans and other mammals since the mammalian array is designed to cover the most conserved regions across different mammalian genomes. To characterize the CpGs that change with macaque age (age-related CpGs) in different tissues, epigenome-wide association studies were carried out, which showed clear tissue-specificity of age-related CpGs (and their proximal genes) (**Figure 4A**). Hence, aging effects in one tissue do not appear to be reflected in another tissue (**Supplementary Figure 3**). This, however, may be owed to the limited sample size in non-blood tissue (**Table 1**).

**Figure 4.**
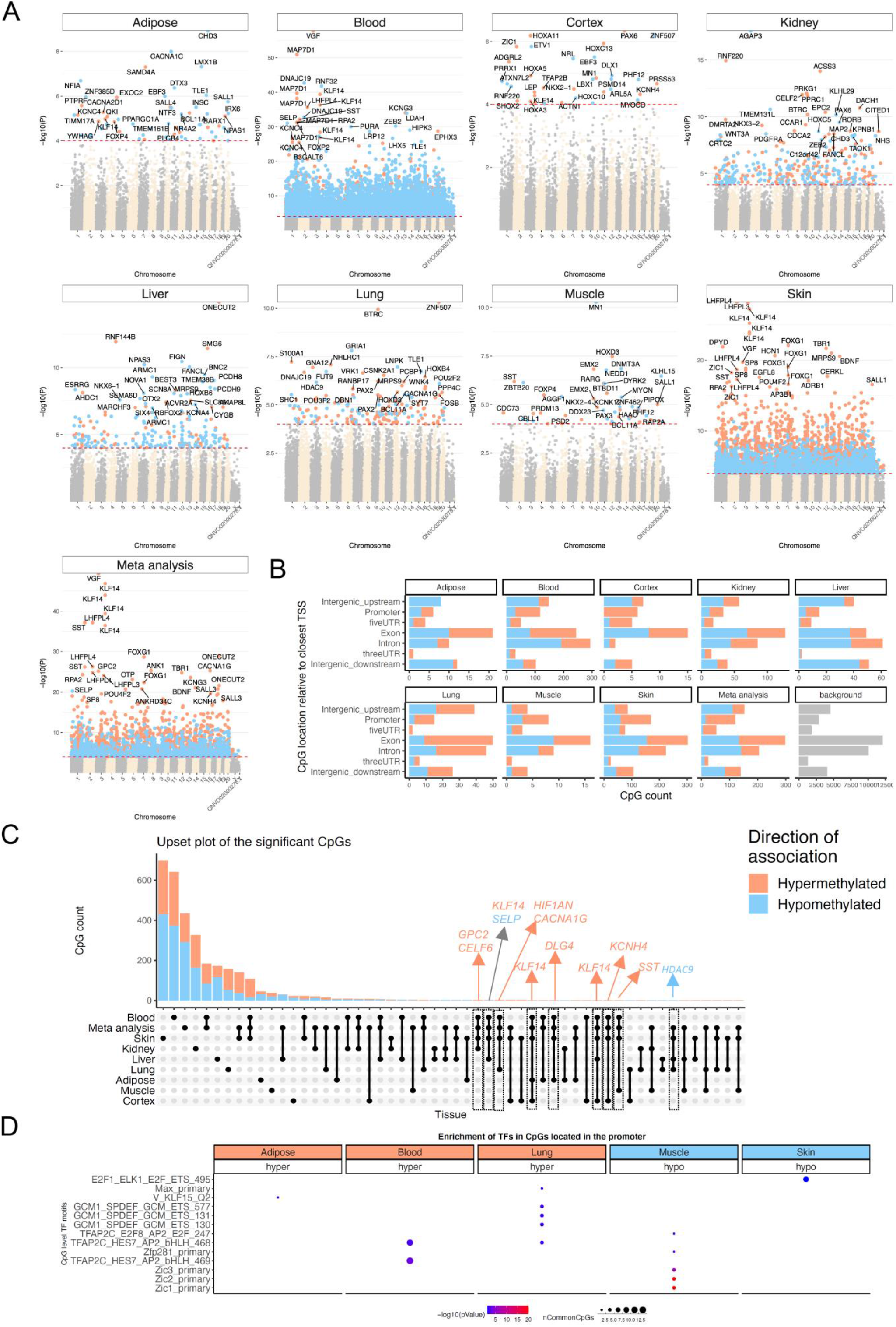
Epigenome-wide association (EWAS) of chronological age in adipose, blood, cerebral cortex, kidney, liver, lung, muscle, and skin of rhesus macaque. A) Manhattan plots of the EWAS of chronological age. The coordinates are estimated based on the alignment of Mammalian array probes to Mmul_10.100 genome assembly. The direction of associations with p < 10^-4^ (red dotted line) is highlighted by red (hypermethylated) and blue (hypomethylated) colors. Top 30 CpGs was labeled by the neighboring genes. B) Location of top CpGs in each tissue relative to the closest transcriptional start site. Top CpGs were selected at p < 10-4 and further filtering based on z score of association with chronological age for up to 500 in a positive or negative direction. The number of selected CpGs: adipose, 62; blood, 1000; cerebral cortex, 40; kidney, 380; liver, 230; lung, 186; muscle, 47; skin, 1000; and meta-analysis, 1000. The grey color in the last panel represents the location of 36733 mammalian BeadChip array probes mapped to Mmul_10.100 genome. C) Upset plot representing the overlap of aging-associated CpGs based on meta-analysis or individual tissues. Neighboring genes of the overlapping CpGs were labeled in the figure. D) Transcriptional motif enrichment for the top CpGs in the promoter and 5’UTR of the neighboring genes. The enrichment was tested using a hypergeometric test (**Methods**).

To identify CpGs whose methylation are most affected by age in all the tissues analyzed, DNAm changes were analyzed at a nominal p value < 10^-4^. The top DNAm changes and their proximal genes in each tissue are as follows: adipose, *CHD3* promoter (Correlation test Z statistic z = −6); blood, *VGF* promoter (z = 16); cerebral cortex, *PAX6* 5’UTR (z = 5); kidney, *AGAP3* intron (z = −8.7); liver, *ONECUT2* exon (z = 7.9); lung, distal intergenic region upstream of *ZNF507* (z = 6.5), and *GRIA1* promoter (z= −5.7); muscle, *MN1* intron (z = −6.5); and skin, *LHFPL4* intron (z = 11). Meta-analysis of these eight tissues, showed the top DNAm changes to include hypermethylation in *VGF* promoter (z = 14.8), four CpGs in *KLF14* promoter (z = 12.7 to 14.5), SST promoter (z = 12.9), and *LHFPL4* exon (z = 12.8) (**Figure 4A**). To identify CpGs that exhibited consistent age-associated methylation change across multiple (but not necessarily all) tissues, we generated an upset plot, which can be interpreted as a generalization of a Venn diagram. The upset plot analysis highlighted four CpGs in the *KLF14* promoter as being age-related in at least 4 tissues (adipose, blood, cortex, and skin, **Figure 4C**). The *KLF14* promoter controls expression of the *KLF14* protein, which is itself a transcriptional factor that regulates the expression of TGFBII receptor.

Age-associated CpGs in different tissues were found to be distributed in genic and intergenic regions that can be defined relative to transcriptional start sites (**Figure 4B**). However, in tissues with sufficient sample numbers (blood and skin), CpGs located in promoters and 5’UTRs had a higher percentage of DNAm change than the background. Moreover, the DNAm changes in promoter and 5’UTR were mainly hypermethylation in all tissues. This result parallels prior observed patterns in DNAm aging in other species. We proceeded to identify putative transcriptional factors whose binding motifs were enriched for the top CpGs located in promoter or 5’UTR with DNAm changes, in either direction and in each tissue (**Figure 4D**). The top TF motifs were Zic1 and Zic2, which had 5 CpGs that become less methylated with age in muscle. These ZIC1 and ZIC2 transcription factors are particularly interesting because they regulate the expression of the *APOE* gene, which is associated with longevity and is the most commonly identified genetic risk factor of Alzheimer’s disease (*24*). Thus, methylation change in this motif might underlie age-associated expression in this protein. For blood and lung, the top enriched motif is the TFAP2C (AP-2 gamma transcriptional factor) binding site that becomes increasingly hypomethylated with age. This motif also exhibited similar age-related changes in other mammalian species and is associated with genes that are involved in cell-cycle arrest, germ cell development, and implicated in several types of cancers (*25, 26*).

## Discussion

Since the inception of the human pan-tissue epigenetic clock in 2013, the field has surged, and more epigenetic clocks have been developed for more applications incorporating different biological parameters that capture a wider range of health effects than mapped in earlier clocks. The human epigenetic clocks have many biomedical applications, including in human clinical trials where they can be used to assess the subjects’ biological age in response to interventions (*5, 27*).

The utility of these human clocks prompted development of similar ones for other mammalian species. Clocks developed for mice are particularly important as they allow modelling of epigenetic age in an animal for which there is good biological understanding (*9, 10, 19–22*). In addition to its impressive legacy in biological sciences, the advantage of a mouse model lies in no small part to its size, which facilitates its maintenance at an affordable cost, even at high numbers. Despite the many advantageous features, there is still a large gap in translating findings to primates. Hence, nonhuman primates play an indispensable role in preclinical investigations of potential interventions that might slow aging. As a case in point, both the National Institute on Aging and the University of Wisconsin have conducted longitudinal studies in rhesus macaques to determine whether the promising anti-aging intervention, caloric restriction, would also apply to primates and hence, more plausibly translate to human aging (*28–30*). Indeed, these studies have yielded valuable information about the role of diet composition, fasting timing, and overall intake on healthspan and lifespan(*29–31*). Despite their importance, such lifespan and healthspan studies in nonhuman primates can exceed the career span of the investigators and are costly. Therefore, the development of suitable biomarkers for biological age promises to greatly reduce the cost and time needed for carrying out such studies and can accelerate our ability to translate interventions.

As indicated, although the pan-tissue human clock can be directly applied to chimpanzees, which diverged from humans approximately 6.3 million years ago, it cannot be applied to any other nonhuman primates since they are more distantly related to humans. Thus, the development of specific epigenetic clocks for other nonhuman primate species is necessary. A critical step that obviates the species barrier was the development of a mammalian DNA methylation array (HorvathMammalMethylChip40) that profiles 36,000 CpGs with flanking DNA sequences that are conserved across multiple mammalian species. This allows DNA methylation profiling of virtually all mammalian species. The rhesus macaque DNA methylation profiles detailed here were derived from eight tissue types and represent the largest dataset to date of single-base resolution methylomes in highly conserved region across multiple tissues and ages.

This successful derivation of the multiple rhesus macaque epigenetic clocks attests, yet again, to the conservation of epigenetic aging mechanisms across the mammalian class. The macaque clock exhibits impressive age correlation with the vervet monkey clock, a species which diverged 12.5 million years ago. Moreover, the evolutionary conservation of epigenetic aging is further exemplified by demonstrating the feasibility of combining methylation profiles of humans and rhesus macaque. These species diverged 29 million years ago yet a single mathematical formula can be applied to generate human-rhesus macaque clocks. This single formula human-macaque clock is equally applicable to both species, and thereby demonstrates conservation of aging mechanisms, which alternatively could be deduced with the existence of multiple individual clocks for other mammals.

The significance of this unification under one formula has far reaching implications which extend beyond its utility in directly translating age-related findings in rhesus macaques to humans. With this tool, one can consider the root contributions to aging as it affirms the increasing evidence that aging is a coordinated biological process, harmonized throughout the body. This ushers in the possibility that when a regulator or coordinator of aging rate is identified, there is potential to modulate it through interventions. As this mechanism is conserved across species, interventions that successfully alter the epigenetic aging rate of rhesus macaques, as measured using the human-rhesus macaque clock, will likely exert similar effects in humans. If validated, this would be a milestone in aging research.

Although genome- and epigenome-wide analyses often yield a large number of potential target genes and pathways related to aging, it is not immediately obvious which ones are actually relevant. Yet, with repeated analyses of age-related CpGs in different species within the mammalian class, the relevant candidates can be identified. Here once again, the advantage of the HorvathMammalMethylChip40 comes to fore. As a case in point, analysis of datasets derived from this array revealed CpGs within the TFAP2 binding site were increasingly unmethylated with age across different mammalian species including rhesus macaque. Additionally, candidates such as Zic1 and Zic2, which did not feature in previously analyzed mammalian species, were uncovered and may indicate species-specific genes related to aging. Evolutionary selection and adaptation would predict a divergence in genes and pathways between species, and this is akin to other biological processes, such as cell cycle regulation, where a basic mechanism is conserved across species, but special additions, deletions, and modifications are identified in only a select species or group.

Just as there are species differences, age-related DNA methylation changes are tissue-specific. Sample size was a limitation of the current study, and thus we can draw only limited conclusions from our data presented here. This notwithstanding, it is interesting to note that CpGs within the KLF14 promoter were consistently altered with age in four tissues (adipose, blood, cerebral cortex, skin). KLF14 is a transcription factor that regulates the TGFBII receptor. This has potential physiological significance because the ligand of this receptor, TGFB, exerts diverse cellular effects including telomere regulation, unfolded protein response, autophagy, DNA repair, cellular senescence and stem cell aging. As a consequence, TGFB signaling is frequently involved in age-related pathologies such as cardiovascular disease, Alzheimer’s disease, and osteoarthritis (*32*).

This is just one example from our extensive analysis of the rhesus epigenome that has broad tissue application and highlights the need for more in-depth empirical investigations to test and reveal the underlying mechanisms of epigenetic aging. Toward this end, the epigenetic clocks will play a pivotal role in uncovering potential candidates, monitoring aging rates, and testing putative aging interventions. The rhesus epigenetic clocks described here would play an inordinately important role in the translation of such interventions to humans.

## Materials and Methods

### Materials

#### Rhesus macaque

In total, we analyzed N=281 rhesus macaque tissue samples from 8 different sources of DNA (**Table 1**). The rhesus monkeys have been housed continuously at the NIH Animal Center, Poolesville, MD. The animal center is fully accredited by the American Association for Accreditation of Laboratory Animal Care, and all procedures were approved by the Animal Care and Use Committee of the NIA Intramural Program. Monkeys were of a heterogenous genetic background, both Chinese and Indian origin.

Monkeys were housed individually in standard nonhuman primate caging on a 12h light/12h dark cycle, room temperature 78+/-2 degrees humidity at 60+/-20%. All monkeys had extensive visual, auditory, and olfactory but limited tactile contact with monkeys housed in the same room. Monkeys received 2 meals per day at estimated ad libitum levels throughout the study. Water was always available ad libitum. Monkeys were monitored minimally 3 times daily by trained animal care staff.

#### Sample Collection

Monkeys were fasted overnight, approximately 16-18 hours. Monkeys were anesthetized with either Ketamine, 7-10 mg/kg, IM or Telazol, 3-5 mg/kg, IM. Blood samples were obtained by venipuncture of the femoral vein using a vacutainer and EDTA tubes. Samples were immediately placed on dry ice and stored at −80 degrees. Skin samples were collected at the same time from an alcohol-wiped area of the back between the shoulder blades. Omental fat, kidney, liver, lung, skeletal muscle, and brain cortex were collected during necropsies scheduled for other study purposes. At that time, tissues were flash frozen in liquid nitrogen following collection and stored at −80 degrees. These tissues were selected for use based on having matching blood samples. None of the monkeys were sacrificed for this study.

#### Vervet monkey animals

The vervet monkey data are described in a companion paper (*33*).

#### Human tissue samples

To build the human-rhesus macaque clock, we analyzed previously generated methylation data from n=850 human tissue samples (adipose, blood, bone marrow, dermis, epidermis, heart, keratinocytes, fibroblasts, kidney, liver, lung, lymph node, muscle, pituitary, skin, spleen) from individuals whose ages ranged from 0 to 93 years. The tissue samples came from three sources: tissue and organ samples from the National NeuroAIDS Tissue Consortium (*34*), blood samples from the Cape Town Adolescent Antiretroviral Cohort study (*35*)., skin and other primary cells provided by Kenneth Raj (*36*). Ethics approval (IRB#15-001454, IRB#16-000471, IRB#18-000315, IRB#16-002028).

#### DNA methylation profiling

We generated DNA methylation data using the custom Illumina chip “HorvathMammalMethylChip40”. By design, the mammalian methylation array facilitates epigenetic studies across mammalian species (including rhesus macaques and humans) due to its very high coverage (over thousand-fold) of highly-conserved CpGs in mammals. Toward this end, bioinformatic sequence analysis was employed to identify 36,000 highly conserved CpGs across 50 mammalian species (Arneson, Ernst, Horvath, in preparation). These 36k CpGs exhibit flanking sequences that are highly conserved across mammals. In addition, the custom array contains two thousand probes selected from human biomarker studies. Each probe is designed to cover a certain subset of species. The particular subset of species for each probe is provided in the chip manifest file and can be found at Gene Expression Omnibus (GEO) at NCBI as platform GPL28271. The SeSaMe normalization method was used to define beta values for each probe (*37*).

#### Penalized Regression models

Details on the clocks (CpGs, genome coordinates) and R software code are provided in the Supplement. Our pan-tissue clock for rhesus macaque is based on 71 CpGs that are present on a custom chip (HorvathMammalMethylChip40). Our human-rhesus macaque epigenetic clock for chronological age is based on 508 CpGs. Another human-rhesus macaque epigenetic clock for relative age is based on 623 CpGs. We developed epigenetic clocks for rhesus macaques by regressing chronological age on the CpGs on the mammalian array. We used all tissues for the pan-tissue clock.

Penalized regression models were created with the R function “glmnet” (*38*). We investigated models produced by both “elastic net” regression (alpha=0.5). The optimal penalty parameters in all cases were determined automatically by using a 10 fold internal cross-validation (cv.glmnet) on the training set. By definition, the alpha value for the elastic net regression was set to 0.5 (midpoint between Ridge and Lasso type regression) and was not optimized for model performance. We performed a cross-validation scheme for arriving at unbiased (or at least less biased) estimates of the accuracy of the different DNAm based age estimators. One type consisted of leaving out a single sample (LOOCV) from the regression, predicting an age for that sample, and iterating over all samples.

For the cross-validation procedure, the penalized regression algorithm automatically selected a different sets of CpGs from the array for each fold. In case of LOO cross validation, the CpG selection was based on n-1 observations. A critical step is the transformation of chronological age (the dependent variable). While no transformation was used for the pan-tissue clock for rhesus macaque, we did use a log linear transformation for the dual species clock of chronological age (Supplementary Methods).

#### Relative age estimation

To introduce biological meaning into age estimates of rhesus macaques and humans that have very different lifespan, as well as to overcome the inevitable skewing due to unequal distribution of data points from rhesus macaques and humans across age range, relative age estimation was made using the formula: Relative age= Age/maxLifespan where the maximum lifespan for rhesus macaques and humans were set to 42 years and 122.5 years, respectively. The maximum lifespan for the two species was chosen from the updated version of the *anAge* data base (*23*).

#### Epigenome wide association studies (EWAS) of age

EWAS was performed in each tissue separately using the R function “standardScreeningNumericTrait” from the “WGCNA” R package (*39*). Next, the results were combined across tissues using Stouffer’s meta-analysis method. Our epigenome wide association test studies of chronological age reveal that aging effects in one tissue are sometimes poorly conserved in another tissue.

### Transcription factor enrichment and chromatin states

The FIMO (Find Individual Motif Occurrences) program scans a set of sequences for matches of known motifs, treating each motif independently (*40*). We ran TF motif (FIMO) scans of all probes on the HorvathMammalMethyl40 chip using motif models from TRANSFAC, UniPROBE, Taipale, Taipaledimer and JASPAR databases. A FIMO scan p-value of 1E-4 was chosen as cutoff (lower FIMO p-values reflect a higher probability for the local DNA sequence matching a given TF motif model). This cutoff implies that we find almost all TF motif matches that could possibly be associated with each site, resulting in an abundance of TF motif matches. We caution the reader that our hypergeometric test enrichment analysis did not adjust for CG content.

### URLs

## Acknowledgements

This study was supported by the Paul G. Allen Frontiers Group (PI SH). The rhesus macaque samples were contributed by the NIA NHP Core which is supported by the Intramural Research Program of the National Institute on Aging, NIH. Human tissue sample collection was supported by NIH funding through the NIMH and NINDS Institutes by the following grants: Manhattan HIV Brain Bank (MHBB): U24MH100931; Texas NeuroAIDS Research Center (TNRC): U24MH100930; National Neurological AIDS Bank (NNAB): U24MH100929; California NeuroAIDS Tissue Network (CNTN): U24MH100928 Data Coordinating Center (DCC): U24MH100925. Human blood samples were supported by R21MH107327. The contents are solely the responsibility of the authors and do not necessarily represent the official view of the NNTC or NIH.

## Authors contributions

The statistical analysis was carried out by JAZ, SH, ATL, AH, CB, JE. Rhesus tissues were provided by the NIA NHP Core and JAM. AJJ contributed vervet tissues. SH, KR, ATL drafted the article. JAM edited the manuscript. The study was conceived of by SH.

## Supplementary Figures

**Supplementary Figure 1.**
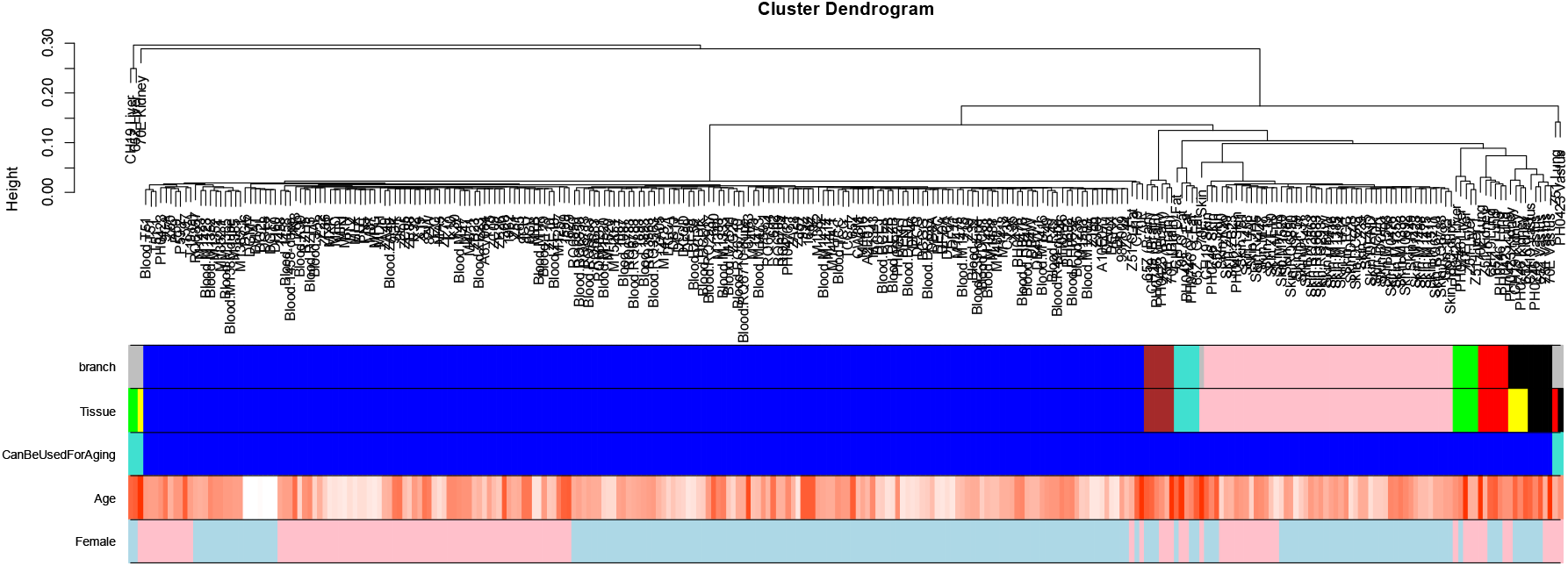
Unsupervised hierarchical clustering of tissue samples. Average linkage hierarchical clustering based on the interarray correlation coefficient (Pearson correlation). A height cut-off of 0.05 led to branch colors that largely correspond to Tissue type (second panel). A handful of arrays are severe outliers (left- and right most clusters) as indicated by the third color band (turquoise samples are severe outliers). These technical outliers probably result from insufficient amounts of DNA. These outlying samples were removed from the analysis.

**Supplementary Figure 2:**
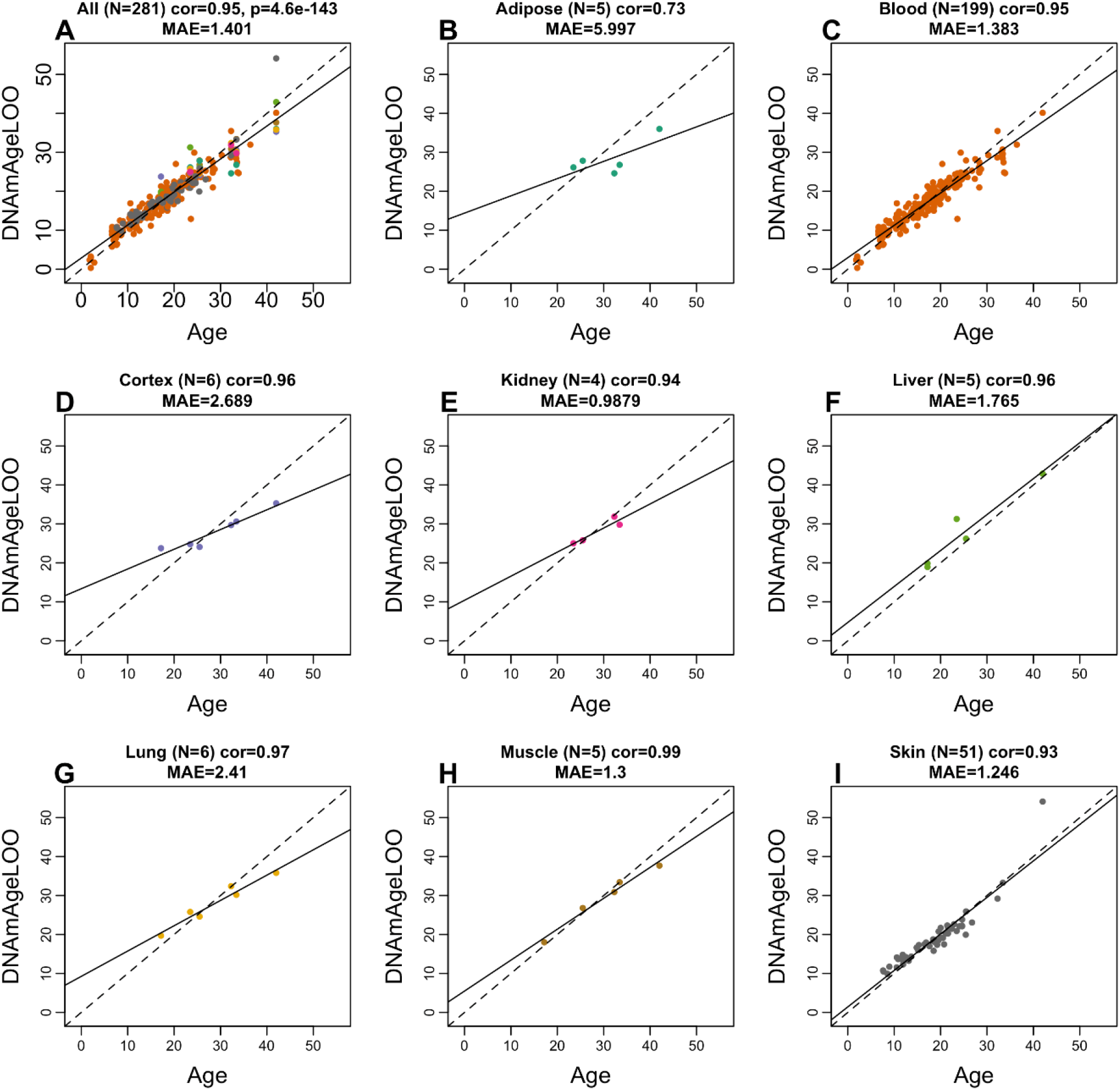
Pan-tissue clock for rhesus macaque applied to different tissues. Each panel correlates the chronological age (x-axis) at the time of tissue collection with the leave-one-out (LOO) estimate of DNA methylation age. A) All tissues. Dots are colored by tissue type as indicated in the remaining panels. Results for B) adipose, C) blood, D) brain cortex, E) kidney, F) liver, G) lung, H) muscle, I) skin. Each panel reports the sample size (N), Pearson correlation, and the median absolute error, i.e. the median of the absolute difference between DNAmAgeLOO and chronological age. The dashed line is the diagonal y=x. The solid line corresponds to the least-squares regression line.

**Supplementary Figure 3.**
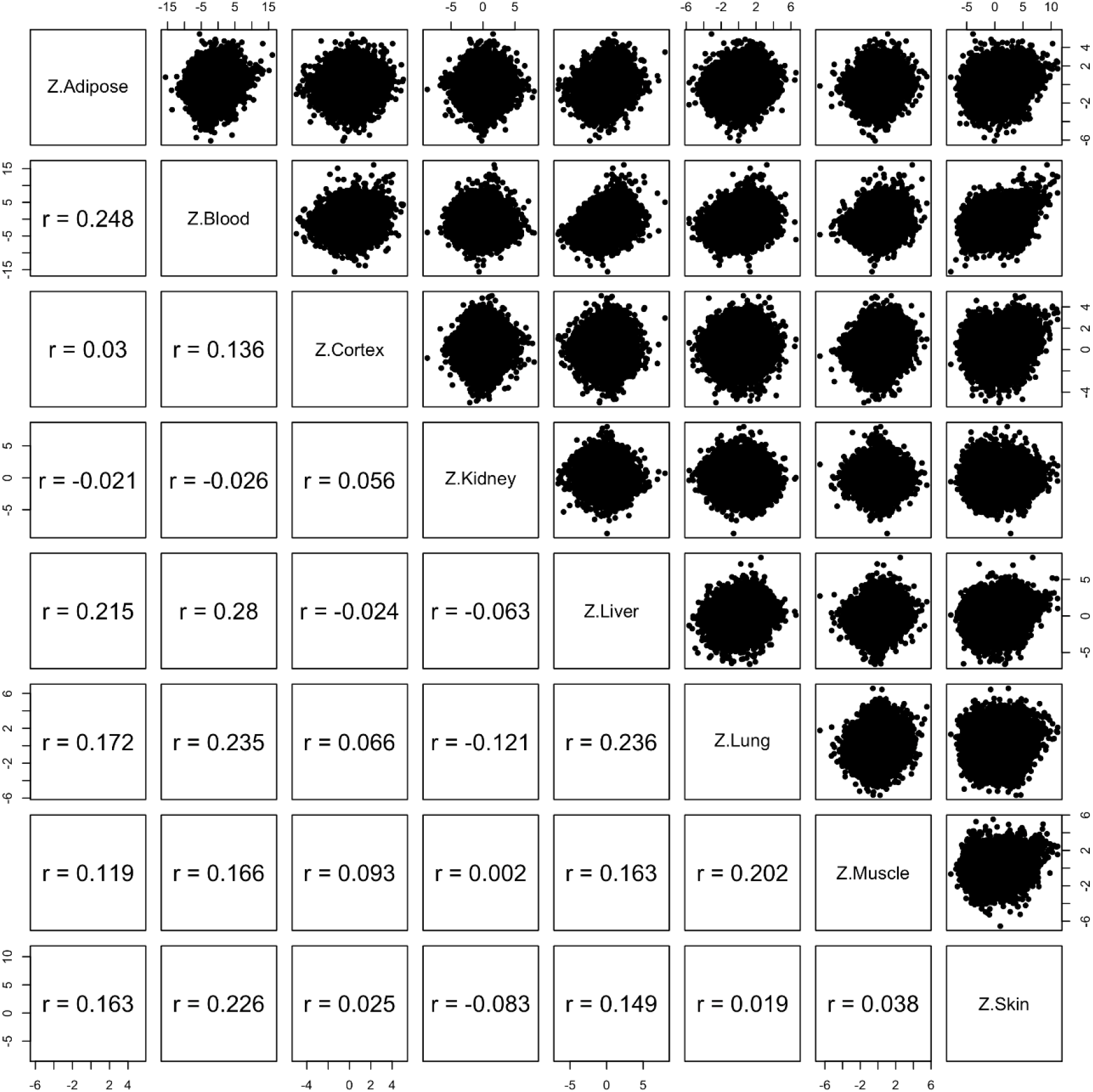
EWAS was performed in each rat tissue separately using the R function “standardScreeningNumericTrait” from the “WGCNA” R package. Z statistics from correlation tests in different tissues. Upper panel report scatter plots for Z statistics in different tissues. Lower panels: corresponding Pearson correlation coefficients.

